## Supplementary figures and images for "Katanin, kinesin-13 and ataxin-2 inhibit premature interaction between maternal and paternal genomes in *C. elegans* zygotes"

### Figure S1

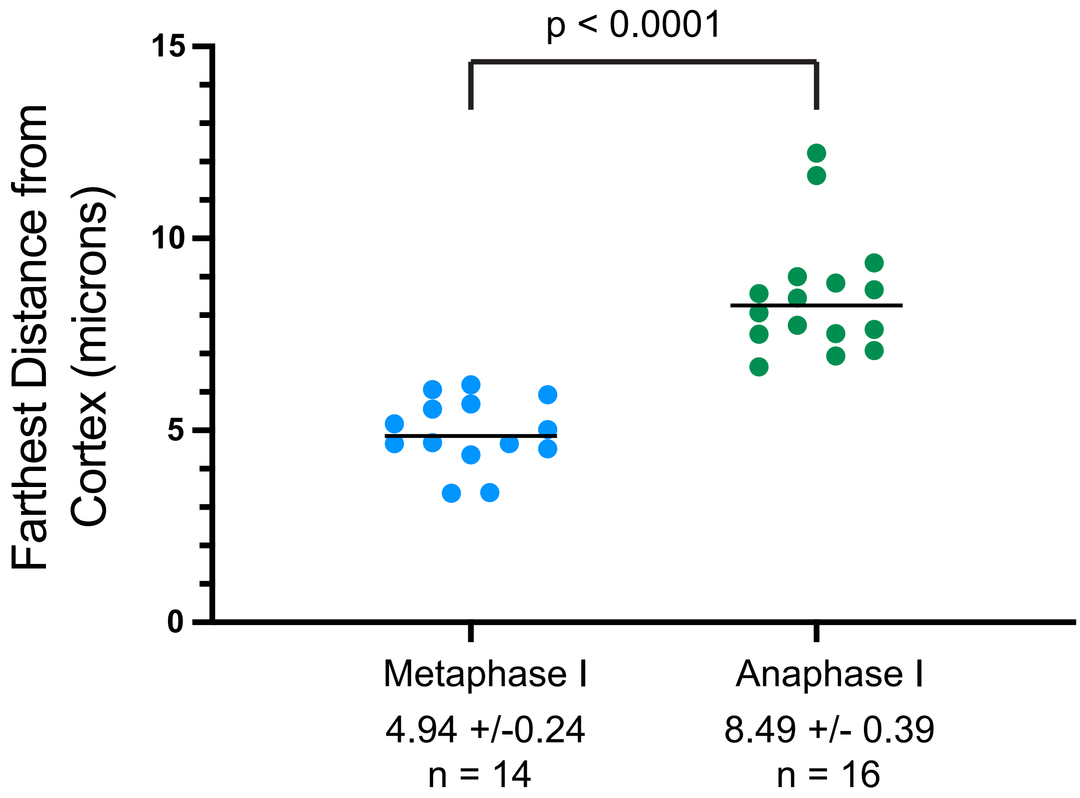

### Figure S2

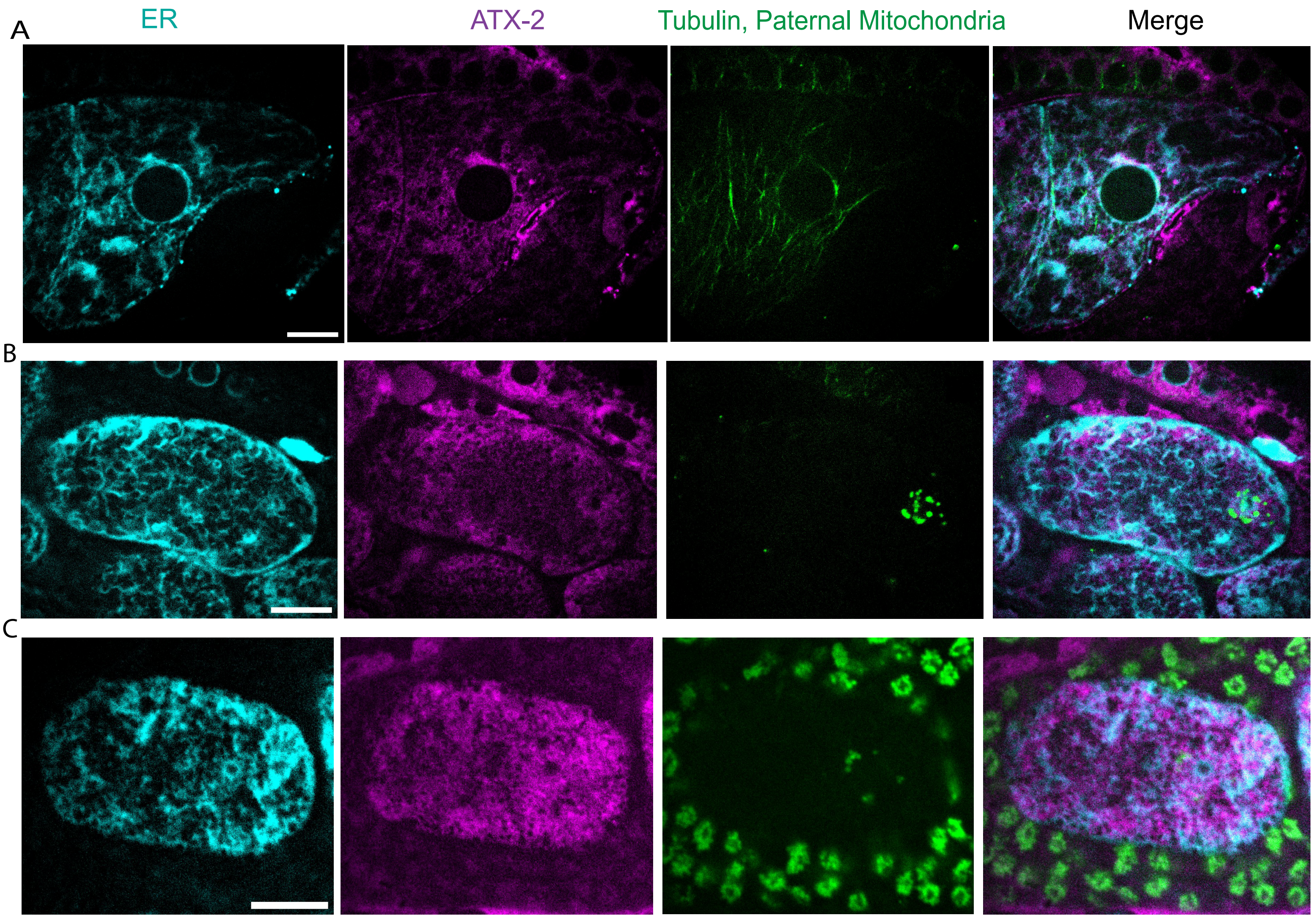

### Figure S3

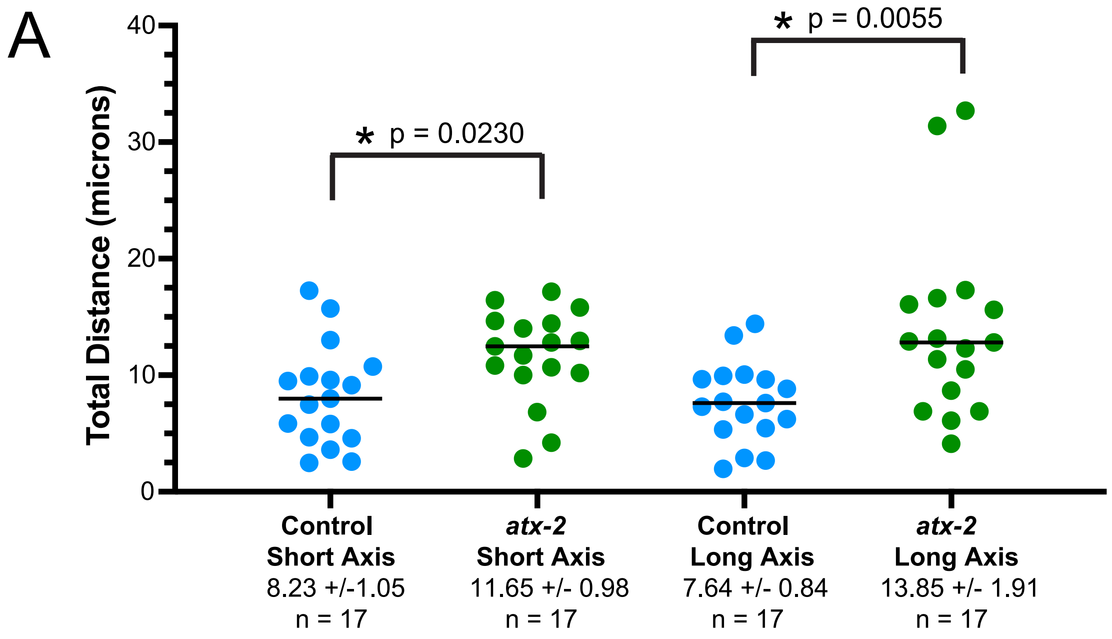
