## Supplementary material for "Katanin, kinesin-13 and ataxin-2 inhibit premature interaction between maternal and paternal genomes in *C. elegans* zygotes": Table S1

**Table S1. *C. elegans* strains**

| Strain Name | Other Name | Genotype Description |
|---|---|---|
| N2 Bristol | | |
| FM111 | WH327 | *unc-119(ed3)* III; *ojls23 [pie-1p::GFP::C34B2.10]* |
| FM302 | CB4108 | *fog-2(q71) V* |
| FM498 | BN580 | *baf-1(bq12[gfp::baf-1]) III* |
| FM500 | | *baf-1(bq12[gfp::baf-1]) III; itIs37 [pie-1p::mCh::H2B::pie-1 3'UTR + unc-119(+)] IV* |
| FM539 | JU2083 | *Caenorhabditis macrosperma* wild isolate |
| FM602 | JJ2586 | cox-4(zu476[cox-4::eGFP::3xFLAG]) I |
| FM638 | | *ojls23 [pie-1p::GFP::C34B2.10]; wjIs76[Cn_unc-119(+); pie-1p::mKate2::tba-2]* |
| FM647 | BCN9071 | *vit-2(crg9070[vit-2::gfp]) X* |
| FM653 | KWN724 | *sdhc-1(jbm1 [sdhc-1::mCherry]) III; him-5(e1490) V* |
| FM727 | | *egxSi126 [mex-5p::hsp-3(aa1-19)::halotag::HDEL::pie-1 3'UTR + unc-119(+)] I; vit-2(crg9070[vit-2::gfp]) X* |
| FM862 | | *atx-2(syb5389; ATX-2::AID::GFP) III; ieSi38 [sun-1p::TIR1::mRuby::sun-1 3'UTR + Cbr-unc-119(+)] IV; wjIs76[Cn_unc-119(+); pie-1p::mKate2::tba-2]* |
| FM932 | | duSi29{pFM1994[TMCO1::GFP(GLO)::SSPB(nanoGLO)]II} *; [pie-1p-mCh::PH(PLC1delta1) + unc-119(+)]V* |
| FM956 | | *ojls23; wjIs76[Cn_unc-119(+); pie-1p::mKate2::tba-2]; atx-2(syb5389; atx-2::AID::GFP); ieSi38 [sun-1p::TIR1::mRuby::sun-1 3'UTR + Cbr-unc-119(+)] IV* |
